# A Control-Theoretic Account of Cognitive Effort in Working Memory

**DOI:** 10.64898/2026.10.11.758303

**Authors:** Tahereh Gholaminejad, Clay B. Holroyd

## Abstract

Cognitive effort is closely linked to prefrontal control, but the control demands imposed by working memory remain poorly understood at the level of neural circuits. We formulated frontoparietal regulation as a feedback-control problem in a conductance-based spiking network comprising an intraparietal ring attractor and a recurrent dorsolateral prefrontal cortex (dlPFC) actuator. A local data-driven task–actuator model was identified around the cue-offset mnemonic state and used to design linear-quadratic regulation. Increasing memory load produced a strong monotonic rise in realized post-cue control effort, driven primarily by sustained dlPFC recruitment. Cue-aligned selective routing supported higher mnemonic loads with substantially less control effort than spatially nonspecific routing. Mean-matched open-loop stimulation reproduced much of the nominal retention benefit, whereas state-dependent feedback provided greater robustness when mnemonic activity was perturbed. Finite-horizon analysis revealed a strongly anisotropic, effectively low-dimensional control geometry dominated by a common mnemonic direction, with much weaker access to item-differentiating directions. These results link increasing working-memory demand to greater sustained prefrontal control requirements and show how frontoparietal routing geometry shapes the efficiency and robustness of mnemonic regulation.

## 1. Introduction

The neural origin and functional purpose of cognitive effort and mental fatigue have been debated for decades [1-2]. Cognitive effort is closely associated with the cognitive-control functions of prefrontal cortex [3-4], yet why prefrontal control should be experienced as effortful remains unclear.

In the influential framework of Miller and Cohen [3], prefrontal activity maintains goal-related representations that bias processing in downstream neural systems. This provides a natural control-theoretic interpretation of prefrontal activity as a control signal that regulates other neural dynamics. Recent control-theoretic accounts make this connection explicit by proposing that cognitive effort is related to the control demands placed on prefrontal systems when regulating downstream neural activity [1]. Within the “controllosphere” account, prolonged prefrontal control activity is hypothesized to promote the accumulation of neural waste products, while the aversive experience of cognitive effort encourages pauses in control that permit recovery [1,5].

A central prediction of this account is that, across cognitive domains, task states associated with greater cognitive effort should impose greater control-energy demands when computational models of these processes are analyzed using control theory. For example, a recent application of LCT to a simple computational model of Stroop-task performance [6] showed that trials with high stimulus–response conflict required more control energy than trials with low stimulus–response conflict [7], consistent with evidence that cognitive conflict is experienced as effortful and carries negative or aversive value [8,9]. Crucially, the theory holds this to be true despite the wide variety of cognitive processes associated with cognitive effort, task designs used to probe those cognitive processes, and computational implementations of the tasks: across domains, application of LCT to these models are predicted to confirm that states associated with cognitive effort demand relatively more control energy. Conversely, the theory predicts that computational models of cognitive effort that do not exhibit this property are incorrect as models of cognitive effort [7]. Here, using control energy as a proxy for cognitive effort in this formal model-based sense, we test this prediction directly in a biophysically grounded model of working memory.

To demonstrate the generality of this approach, here we apply LCT to a different, effortful cognitive process: working memory. Working memory is a limited-capacity system that temporarily maintains and manipulates information needed for ongoing cognition and goal-directed behaviour [10,11]. Importantly for the present account, increasing working-memory demand increases the subjective cost of cognitive engagement [12], and working memory is strongly associated with cognitive fatigue [13]. Furthermore, visual working memory (VWM) tasks provide a particularly useful setting in which to examine the limits and control demands of short-term maintenance [14]. VWM maintains information after the corresponding sensory input is no longer available, but only a limited number of representations can be retained reliably. Classical change-detection studies place this limit at approximately four simple objects [15], although performance varies substantially across individuals and trials as a function of attentional control [16]. These observations suggest that working memory capacity is not determined solely by storage limitations but is also shaped by the control processes that support ongoing maintenance. Together, these features make VWM a suitable domain in which to test whether greater task demand is accompanied by greater formal control demand.

At the circuit level, working memory maintenance has long been associated with stimulus-selective persistent activity [17], although recent population recordings indicate that mnemonic information need not be expressed continuously in single-trial spiking [18]. Recurrent attractor models nevertheless provide a well-established framework for studying the circuit mechanisms that stabilize memory representations. In the spatial working memory model of Compte et al. [19], recurrent excitation, mediated primarily by NMDA receptors and balanced by strong recurrent inhibition, sustains localized activity after cue offset.

Here we apply LCT to a computational model of VWM by Edin et al. [20], who extended this framework to working memory capacity by showing that lateral inhibition between simultaneously maintained representations limits the number of stable mnemonic activity bumps in IPS. They proposed that additional excitatory input from dlPFC can counteract this inhibition at high memory loads and thereby increase parietal capacity, a prediction supported by their analysis of frontoparietal activity during high-load performance. This architecture provides a natural test bed for the controllosphere account: the IPS network supplies nonlinear mnemonic dynamics, whereas dlPFC provides an anatomically distinct top-down pathway capable of regulating those dynamics. Our primary question is whether increasing working memory load places greater control-energy demands on the dlPFC pathway, as predicted by the controllosphere account of cognitive effort.

A second question concerns how this top-down support is spatially organized. Edin et al. [20] modelled dlPFC support as spatially nonspecific excitation of the parietal network, whereas physiological studies indicate that frontoparietal communication can also convey spatially and content-specific information [21–24]. We therefore compare Edin-like spatially uniform input with a cue-aligned selective routing profile while matching their mean dlPFC→IPS projection strength. This comparison asks whether the geometry of top-down access, rather than simply the overall magnitude of prefrontal recruitment, can alter effective VWM capacity. Given the substantial variation in VWM performance across individuals, including variation associated with attentional control [16], differences in effective frontoparietal routing could represent one potential contributor to such variability, although the present model does not attempt to model individual differences directly. This comparison also allows us to ask whether the spatial organization of prefrontal access changes the control energy required to support a given memory load.

Beyond the primary question of cognitive effort, this formulation allows us to characterize how the geometry of prefrontal access constrains the control energy required to regulate mnemonic states. Control theory has increasingly been used to address related questions in neural systems. At the whole-brain level, network-control approaches have examined how structural organization constrains state transitions, controllability, and control energy [25–31]. Related formulations have also been applied to local neural circuits and recurrent networks, including optimal control of motor-preparatory dynamics and spiking neural networks [32–34]. Working-memory studies provide particularly relevant precedents. Braun et al. [35] found that a more demanding 2-back state was less stable and required greater control energy than a lower-demand control state, while Cai et al. [36] reported load-dependent changes in functional controllability and asymmetric control roles across distributed salience and frontoparietal circuits. Together, this work provides a formal framework for asking not only whether a neural pathway can alter network activity, but also which state-space directions it can influence and how much control effort those transitions require.

These questions depend critically on the geometry of neural population activity. Experimental and theoretical work indicates that neural trajectories occupy structured, often low-dimensional regions of population state space [37–40]. For example, Sadtler et al. [41] showed that monkeys learned brain–computer-interface mappings that remained within the intrinsic manifold of motor-cortical activity more readily than mappings that required activity outside it. Although mnemonic regulation poses a different problem, the same geometric principle is relevant here: the effectiveness and energetic demands of dlPFC control should depend on how the available input pathway is situated relative to task-relevant directions in IPS state space. This motivates a controllability analysis of which mnemonic directions are robustly accessible through the dlPFC→IPS pathway and how the geometry of that access shapes the control energy required for regulation.

Applying LCT to this system poses an important methodological challenge: the controlled plant is a high-dimensional, nonlinear, and stochastic spiking network, whereas LQR requires a tractable local state-space model. Rather than replacing the spiking circuit with a globally valid linear or mean-field surrogate, we retain the full conductance-based network as the controlled plant and identify its local cue-defined task-space dynamics directly from perturbation responses of the spiking system. Because the command reaches IPS indirectly through recurrent dlPFC activity and dlPFC→IPS synaptic dynamics, we additionally identify a low-dimensional representation of the actuator pathway and combine it with the mnemonic task state for controller design.

Using the identified local task–actuator model, we design discrete-time LQR control while retaining the nonlinear spiking network as the controlled plant. Our primary question is whether increasing working memory load imposes greater model-based control-effort demands on the dlPFC pathway, as predicted by the controllosphere account of cognitive effort. We then examine how cue-aligned routing and finite-horizon task-space access shape this demand. Together, these analyses provide a circuit-level test of whether increasing cognitive demand is accompanied by increasing formal control demands on prefrontal regulation.

## 2. Methods

### 2.1 The parietal spiking plant

The IPS plant is a conductance-based spiking ring attractor containing *N*_*E*_ *=* 1024 excitatory pyramidal neurons and *N*_*I*_ *=* 256 inhibitory interneurons. The IPS network follows the working-memory model of Edin et al. [20], which builds on the conductance-based ring-attractor framework introduced by Compte et al. [19]. Neurons are assigned uniformly spaced preferred angles on a circular domain. Spatially structured recurrent projection weights are defined by the circular Gaussian kernel in Eq. (1),

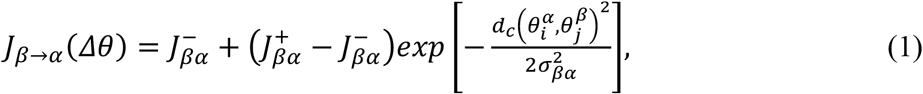

where, *α* and *β* denote the postsynaptic and presynaptic populations, respectively; 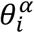and 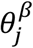are the preferred angles of postsynaptic neuron *i* and presynaptic neuron 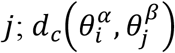is the shortest circular distance; σ_*β α*_is the projection width; and 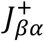and 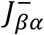 are the peak and baseline weights. The baseline weight is chosen so that the mean incoming weight equals one,

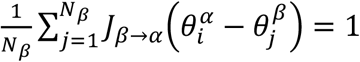

where, *N*_*β*_ is the size of the presynaptic population. The membrane dynamics of each IPS neuron are then described by the conductance-based leaky integrate-and-fire equation,

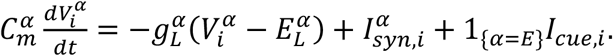

where, 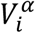is the membrane potential of neuron *i*, 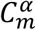is its membrane capacitance, 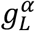is the leak conductance, 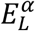is the leak reversal potential, 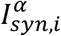 is the total synaptic current, *I*_*cue,i*_ is the cue current, and *1*_*{α=E}*_ restricts the cue to excitatory IPS neurons. Excitatory neurons use 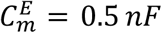, and 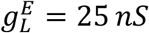 inhibitory neurons use 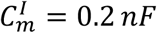, and 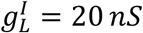. Both populations use the membrane thresholds, resets, and reversal potentials specified by the underlying spiking model [19,20]. The total synaptic current is written in conductance form,

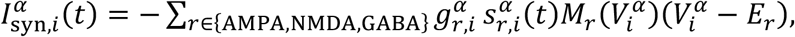

where, *r* indexes receptor type, 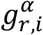 is the corresponding maximal conductance, 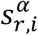is the aggregate gating variable arriving at neuron *i, E*_*r*_ is the receptor reversal potential, and *M*_*r*_*(V)* is the voltage-dependent factor, which equals 1 for AMPA and GABA, whereas NMDA uses a nontrivial magnesium-block factor. Background afferents and dlPFC spikes enter IPS through AMPA conductance, recurrent excitation is NMDA-mediated, and recurrent inhibition is GABA-mediated [19,20]. Importantly, the control command is never injected directly into the IPS membrane equation; all top-down control reaches IPS through dlPFC spiking and the dlPFC→IPS synaptic projection.

AMPA and GABA gating variables undergo spike-triggered increments followed by exponential decay [19,42],

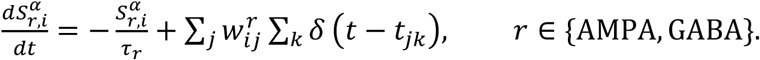

Here, *τ*_*r*_ is the receptor decay time constant, 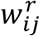is the synaptic increment from presynaptic neuron *j* to neuron *i, δ(⋅)* is the Dirac delta function, and *t*_*jk*_ is the time of the *k*-th spike emitted by neuron *j*. The slower NMDA conductance follows the two-stage kinetic scheme [19],

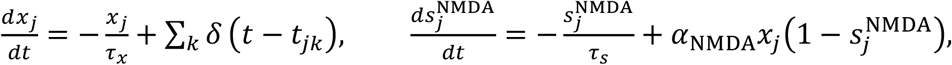

where, *x*_*j*_ is the fast NMDA rise variable, 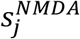 is the slower NMDA gating variable, *τ*_*x*_ and *τ*_*s*_ are their time constants, and *α*_*NMDA*_ is the opening-rate constant. The simulations use *τ*_*AMPA*_ *=* 2 *ms, τ*_*GABA*_ *=* 10 *ms, τ*_*x*_ *= 2 ms, τ*_*s*_ *=* 100 *ms*, and *α*_*NMDA*_ *= 0*.*5 ms*^−1^ . Each IPS neuron receives independent external Poisson AMPA input at a total rate of 1800 Hz per cell. For memory load *L*, cue locations are equally spaced around the ring and are presented from *t =* 0 to *t*_*c*_ *=* 500 *ms* with circular-Gaussian width σ_*C*_ *= 9*.*4*^*°*^ and peak current 0.025 *nA*. After cue offset, the network evolves through a 2500-ms delay period, giving a total trial duration of 3000 ms. Spiking dynamics are simulated in Brian 2 [43] using the integration time step Δ*t =* 0.02 *ms*; while the identified control model and controller operate at their coarser sampling interval Δ*t*_*c*_ *=* 10 *ms*.

### 2.2 Task-space state representation

Controller synthesis and state feedback are formulated in a load-dimensional task space derived from the excitatory IPS NMDA state, rather than in the full spiking-network state space. We use the excitatory NMDA gating state because it is the slow recurrent variable supporting persistent mnemonic activity and provides a smooth representation of the maintained network state at the controller sampling timescale. For each of the *L* cued locations, we define a positive circular-Gaussian direction over the excitatory IPS ring using the same spatial width as the sensory cue. These directions depend only on the cued item locations and are normalized before construction of the task-space readout, defined as follows,

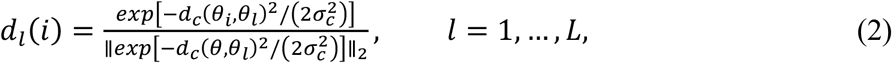

where, *d*_*l*_ is the unit-norm task direction for item *l, θ*_*i*_ is the preferred angle of excitatory IPS neuron *i, θ*_*l*_ is the centre of the *l*-th cued location, σ_*c*_ is the circular-Gaussian width of the sensory cue, *d*_*C*_(·) is the shortest circular distance, and ‖ *·* ‖_2_ denotes the Euclidean norm. The normalized cue-centered directions are stacked as the rows of

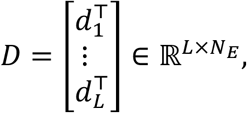

and symmetrically whitened to obtain the orthonormal task-space readout *C* as follows,

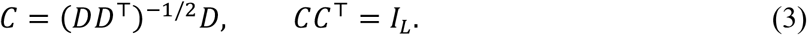

where, *I*_*L*_ is the identity matrix of dimension *L*. Symmetric whitening preserves the row space of *D* unchanged, so the readout *C* spans the same cue-defined subspace while supplying an orthonormal coordinate system for identification, control, and validation at a given load.

At controller sampling time *t*_*k*_ *= kΔt*_*C*_, the full excitatory IPS NMDA gating state vector 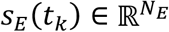 is projected onto the cue-defined task subspace using the readout matrix *C*, yielding the *L*-dimensional task state

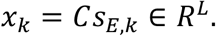

The components of *x*_*k*_ describe the network state within the cue-defined mnemonic subspace. The mnemonic reference is defined directly from the IPS state at cue offset,

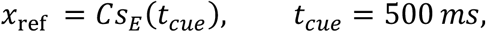

where, *x*_ref_ is the mnemonic reference, *t*_*cue*_ is the cue-offset time, and *s*_*E*_*(t*_*cue*_) is the excitatory IPS NMDA state at cue offset in the trajectory under analysis. Across the compared conditions, the same cue-offset IPS state is used as the initial mnemonic state, ensuring that the delay period begins from an identical sensory-evoked configuration. The reference *x*_ref_ is held fixed throughout the subsequent 2500-ms delay. It is therefore a fixed regulation reference defined by the mnemonic state, not a fitted equilibrium and not an assumption that the uncontrolled IPS dynamics are stationary at cue offset.

For each memory load *L*, the same task-space readout *C* is used for both routing configurations (see Section 2.4), ensuring that flat and selective routing are evaluated within an identical cue-defined mnemonic representation. Differences between the two conditions therefore reflect the physical dlPFC→IPS routing profile and the corresponding controller. The reduced *L*-dimensional state is used for system identification and controller design, while retention outcomes, including the number of surviving mnemonic bumps, are evaluated from the full spiking-network activity.

### 2.3 Equation-free local linearization in task space

The full spiking network defines the plant dynamics, while controller synthesis and controllability analysis use a tractable local discrete-time model in the cue-defined task space. We estimate the intrinsic IPS task-space dynamics using the equation-free coarse time-stepper framework introduced by Kevrekidis et al. [44]. In this framework, short simulations of the microscopic system are used to evaluate the evolution of prescribed coarse variables over a finite time interval. Related equation-free formulations have been applied to neural-network dynamics and to controller design for microscopic and stochastic simulation models [45–47]. Here, the coarse variables are the *L*-dimensional task coordinates defined in Section 2.2, and the coarse time step is the controller sampling interval, Δ*t*_*c*_ *=* 10ms. Identification is performed offline for each memory load *L* and routing configuration.

The use of equation-free identification was motivated by an initial attempt to construct a conventional mean-field reduction. Although such a reduction provides a lower-dimensional description of population activity, in our preliminary analyses it did not capture the local dynamics and spatial bump structure with sufficient fidelity for controller identification. We therefore estimated the local coarse dynamics directly from the spiking network. For each load, we identify a local transition matrix *A*_*x*_ ∈ ℝ^*L×L*^ describing the one-step evolution of small task-state displacements,

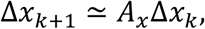

where Δ*x*_*k*_ ∈ ℝ^*L*^ is a local task-space displacement at controller sample *k*. Thus, *A*_*x*_ characterizes the intrinsic evolution of mnemonic-state displacements over one controller interval.

The task-space readout *C* defined in Eq. (3) serves as the restriction operator, with *x = Cs*_*E*_. Local displacements are introduced directly in the excitatory IPS NMDA state. Let 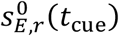 denote the cue-offset NMDA state for identification sample *r*. For each cue-centred direction *d*_*j*_, *j =* 1, *…,L*, defined in Eq. (2), we construct a nearby microscopic state and its corresponding task-space displacement as

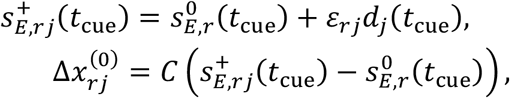

where 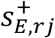 is the displaced excitatory IPS NMDA state, *ε*_*rj*_ *>* 0 is the applied displacement magnitude, *r* indexes the identification samples, and *j =* 1, *…,L* indexes the cue-centred directions. This construction provides the local microscopic initialization, corresponding to the lifting step of the coarse time-stepper.

For each direction *d*_*j*_, the displacement magnitude *ε*_*rj*_ was scaled to the local unforced 10-ms variation of the IPS NMDA state and reduced when necessary to keep the NMDA gating variables within their admissible range.

Starting from the displaced initial NMDA state 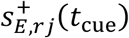 and the corresponding reference initial state 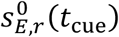, the full conductance-based spiking network is evolved for one controller interval, Δ*t*_*c*_ *=* 10 ms. The resulting excitatory IPS NMDA states at *t*_cue_ *+ Δt*_*c*_ are denoted by 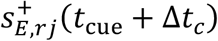 and 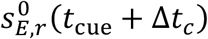 respectively. After restriction through *C*, the one-step task-space displacement is

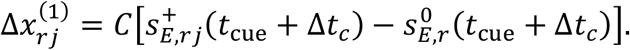

For each identification sample *r*, the displaced and reference trajectories are propagated using the same stochastic input sequence. This common-random-numbers procedure reduces stochastic variability shared by the two trajectories [48]. The pair 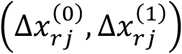 therefore provides one local one-step sample of the mapping from the imposed task-space displacement 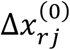 to its evolved value 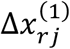. Because only positive NMDA displacements, *ε*_*rj*_ *> 0*, are applied along *d*_*j*_, these samples form a one-sided finite-difference estimate of the local coarse time-stepper. The transition matrix *Â*_*x*_ is estimated by pooling the finite-difference samples across task directions and the identification set,

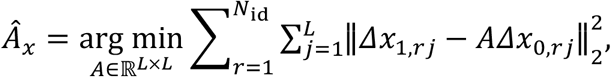

where *N*_id_ is the number of identification samples and *Â*_*x*_ is the least-squares estimate of the local task-space transition matrix. Collecting the initial and one-step displacements as columns of *X*_id_ and *Y*_id_, respectively, gives

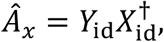

where 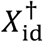 denotes the Moore-Penrose pseudoinverse [49].

Figure 1 summarizes the equation-free identification procedure. The nominal and displaced IPS states are advanced through the original spiking network under matched stochastic input and then restricted to task space to form paired one-step displacements. Pooling these pairs provides the local operator *Â*_*x*_ required for controller synthesis without requiring an explicit closed-form population model. The identified operator *Â*_*x*_ is evaluated using an independent set of composite displacements formed from novel combinations of the cue-defined directions. For each composite displacement, the one-step task-space response measured from the full spiking network is compared with the prediction *Â*_*x*_ Δ*x*^(0)^. This evaluates the generalization of the local linear model from the individual directions used for identification to previously unseen combinations of mnemonic-state displacements. Numerical validation results are reported in Section 3.1.

**Fig. 1.**
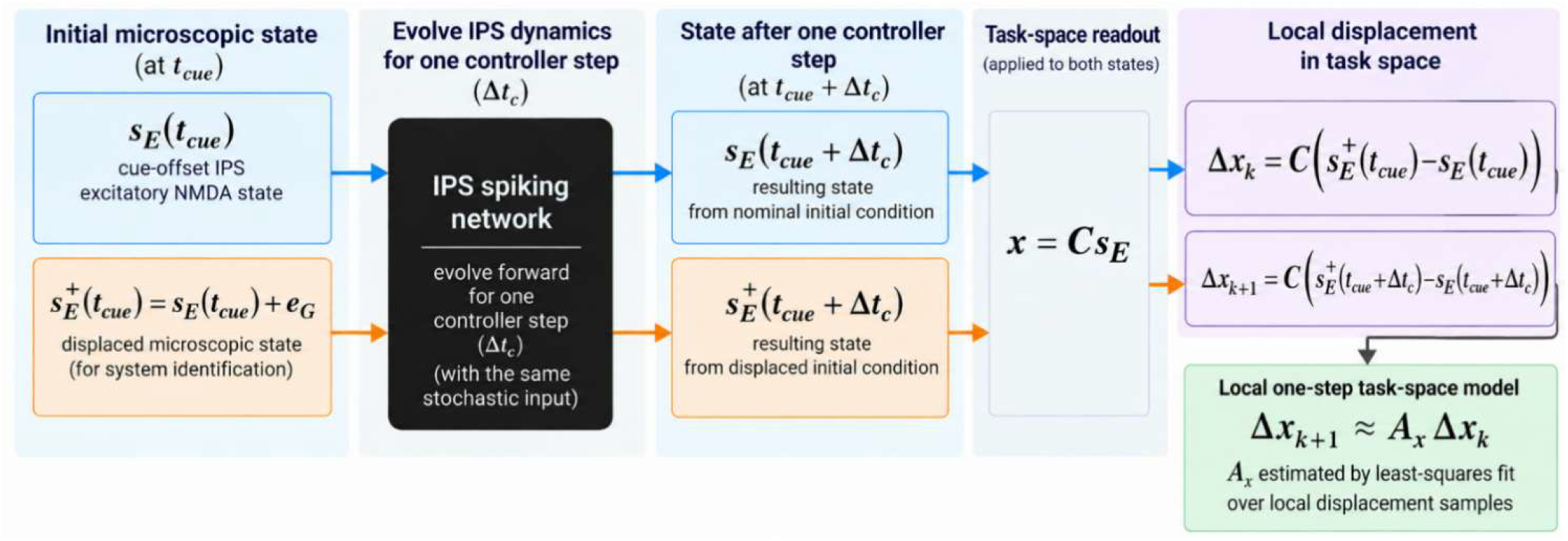
Equation-free identification of the intrinsic IPS task-space dynamics. A small displacement is applied to the cue-offset excitatory IPS NMDA state, and the nominal and displaced microscopic states are evolved for one controller interval under identical stochastic input. Restriction through the cue-defined readout *C* yields paired task-space displacements Δ*x*_*k*_ and Δ*x*_*k*_.

The matrix *Â*_*x*_ supplies the intrinsic IPS task-space transition block of the augmented task-actuator model introduced in Section 2.5.

### 2.4 The prefrontal actuator and spatial routing

The control actuator is an untuned recurrent spiking dlPFC network comprising excitatory and inhibitory neurons, with no spatial structure in its recurrent connectivity. The intrinsic cellular, recurrent-synaptic, and background-input parameters of the spiking dlPFC network are retained from the source model [20] and are held fixed during system identification and controller design. The study-specific components are the external control command and the flat versus cue-aligned dlPFC→IPS routing manipulation described below.

Working-memory retention is first characterized without prefrontal recruitment or post-cue control. To reproduce the prefrontal recruitment manipulation of Edin et al. [20], a 0.15-nA current pulse is applied to dlPFC excitatory neurons during the first 100 ms of cue presentation, establishing persistent prefrontal activity. The same recruitment pulse defines the prefrontal operating state for the feedback-control and mean-matched open-loop conditions. The post-cue control command, Δ*u*_*k*_, is defined as an additional current relative to this recruited state and is applied exclusively to dlPFC excitatory neurons. Its influence on IPS is mediated through dlPFC spiking and the dlPFC→IPS synaptic projection.

Long-range dlPFC→IPS excitation is mediated by AMPA synapses from dlPFC excitatory neurons to IPS excitatory neurons, with a maximal synaptic conductance of 9 nS. In the configuration used here, the corresponding dlPFC→IPS conductance onto IPS inhibitory neurons is zero.

To examine how the spatial organization of prefrontal input affects mnemonic regulation, we compare two dlPFC→IPS routing configurations with matched mean projection strength. In the flat configuration, corresponding to the spatially nonspecific dlPFC→IPS projection in the original Edin et al. model [20], dlPFC excitatory output is distributed uniformly across the IPS excitatory population. In the selective configuration, the same scalar prefrontal output is weighted by a prescribed cue-aligned spatial profile constructed from the cued item locations. The selective routing profile is defined in Eq. (4).

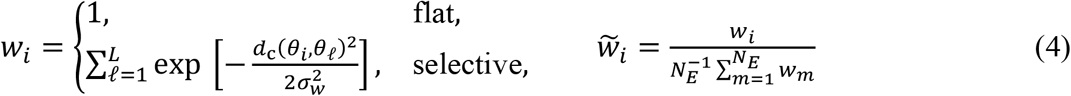

Here, *w*_*i*_ denotes the unnormalized spatial support at excitatory IPS neuron *i*, 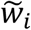 is the corresponding profile normalized to unit mean across the excitatory IPS population, *θ*_*i*_ is the preferred angle of neuron *i, θ*_*l*_ is the centre of cued item *l, L* is the memory load, and σ_*w*_ is the cue-aligned routing width. Unit-mean normalization ensures that flat and selective routing have the same population-averaged dlPFC→IPS projection strength. The selective profile therefore changes the spatial distribution of the scalar prefrontal drive without introducing separate control commands for individual items. The two routing configurations and their relationship to the IPS ring attractor are illustrated schematically in Fig. 2.

**Fig. 2.**
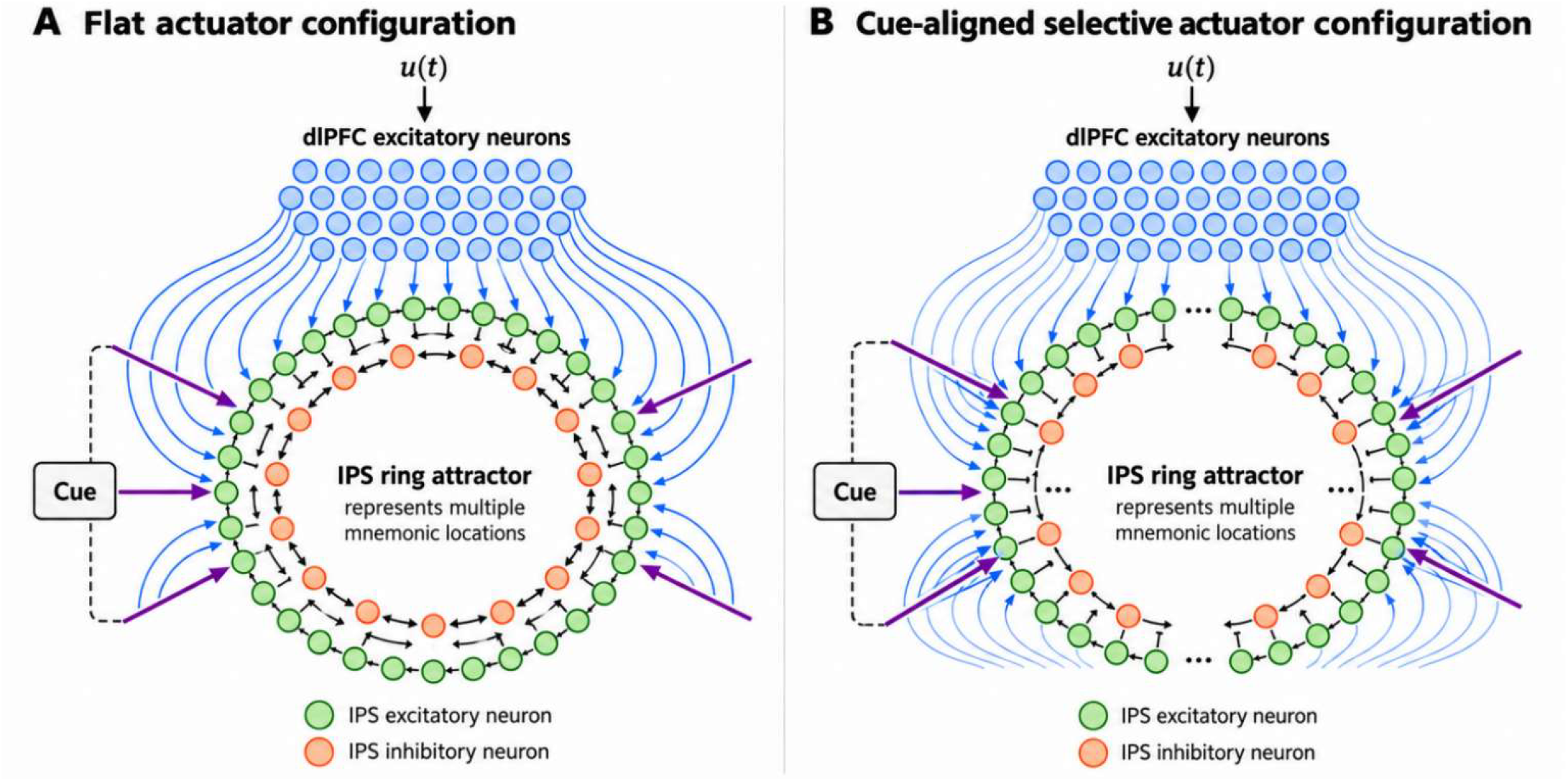
IPS ring attractor and flat versus cue-aligned dlPFC→IPS routing. The selective Gaussian profile is normalized to the same mean projection strength as the flat profile; the scalar post-cue control command is applied through the dlPFC pathway.

### 2.5 Actuator identification and augmented task-actuator model

Because the control command Δ*u*_*k*_ reaches IPS through the recurrent dlPFC network and the intervening dlPFC→IPS synaptic dynamics, its effect on the mnemonic task state is dynamic rather than instantaneous. We therefore augment the task state with two low-dimensional coordinates of the actuation pathway. The first, *p*_*k*_, is the mean excitatory dlPFC NMDA gating variable. The second, *a*_*k*_, is the projection coefficient of the dlPFC-evoked IPS AMPA gating-state vector onto the normalized routing profile defined in Eq. (4).

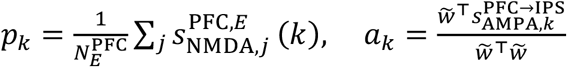

where 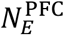 denote the number of excitatory dlPFC neurons, 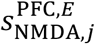 the vector of NMDA gating variables across excitatory dlPFC neurons, and 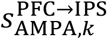 the vector of dlPFC-evoked AMPA gating variables across excitatory IPS neurons at sample *k*. Together, these variables define the fixed two-dimensional actuator state

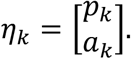

The coordinate *p*_*k*_ captures recurrent dlPFC activation, whereas *a*_*k*_ captures AMPA-mediated transmission from dlPFC to IPS. These actuator coordinates are distinct from the recurrent IPS synaptic gating variables used in the parietal plant. The same actuator-state definition is used for all memory loads and both routing configurations.

For each memory load and routing configuration, the remaining task–actuator dynamics are identified from command-evoked responses of the full spiking network, with *A* fixed at the value obtained in Section 2.3. Short pulse, alternating, and step command sequences are applied to dlPFC, and each commanded trajectory is paired with a matched trajectory having zero post-cue command. At each controller sample, the differences between the paired trajectories define the command-induced task-state deviation Δ*x*_*k*_ and actuator-state deviation Δ*η*_*k*_ *= [Δp*_*k*_, Δ*a*_*k*_]^*T*^.

These measured deviations are used to identify the one-step model

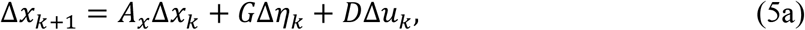

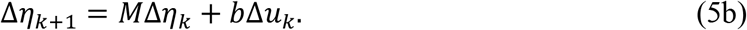

where, Δ*x*_*k*_ is the task-state deviation between the commanded trajectory and its matched zero-command trajectory, Δ*η*_*k*_ is the corresponding actuator-state deviation, and Δ*u*_*k*_ is the scalar post-cue command applied to dlPFC. Because *A*_*x*_ ∈ ℝ^*L×L*^ is fixed, *G* ∈ ℝ^*L×2*^ and *D* ∈ ℝ^*L×1*^ are estimated jointly by least-squares regression of the residual task response Δ*x*_*k+1*_ − *A*_*x*_ Δ*x*_*k*_ onto the predictors Δ*η*_*k*_ and Δ*u*_*k*_. The actuator matrices *M* ∈ ℝ^2 *×2*^ and *b* ∈ ℝ^2 *×1*^ are estimated independently by regressing Δ*η*_*k+1*_ onto the same predictors. Data from all pulse, alternating, and step command experiments are combined when fitting these matrices by least squares.

With *A*_*x*_, *G, D, M*, and *b* determined, Eqs. (5a)–(5b) can be combined into a single augmented state-space representation. We define the augmented state as

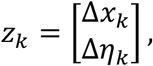

which gives

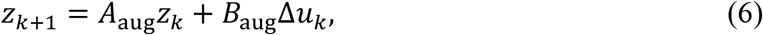

with

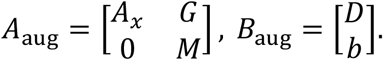

Here, *z*_*k*_ ∈ ℝ^*L+2*^ contains the *L*-dimensional task-state deviation and the two-dimensional actuator-state deviation. The zero lower-left block reflects the implemented network architecture, in which IPS does not project back to dlPFC.

After parameter estimation, the identified augmented model is evaluated on an independent 500-ms multilevel command sequence that is not used for fitting. At each controller sample, the measured augmented state *z*_*k*_ from the full spiking network and the applied command Δ*u*_*k*_ are used to generate the one-step prediction

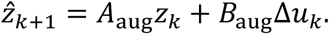

Thus, each prediction spans a single 10-ms controller interval and is initialized from the measured network state at sample *k*.

Prediction accuracy is quantified for each of the *L* task coordinates and the two actuator coordinates, *p* and *a*. Prediction-error magnitude is measured using the normalized root-mean-square error (NRMSE):

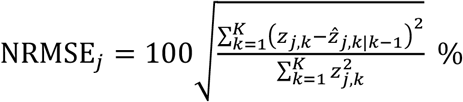

where, *j* indexes the task and actuator coordinates, *K* is the number of validation samples, *z*_*j,k*_ is the measured value of coordinate *j* at sample *k*, and 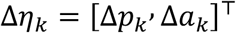 is the corresponding one-step prediction. Normalization by the RMS amplitude of the measured trajectory allows prediction errors to be compared across coordinates with different numerical scales.

Directional agreement between the predicted and measured trajectories is quantified using cosine similarity:

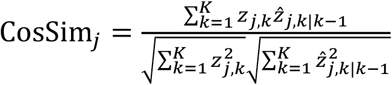

where, CosSim_*j*_ is the cosine similarity between the measured and predicted trajectories of coordinate *j* across the *K* validation samples. Values approaching one indicate close directional agreement between the predicted and measured trajectories. Thus, NRMSE quantifies prediction-error magnitude, whereas cosine similarity quantifies directional agreement.

The validation procedure is performed separately for every memory load and routing configuration. Aggregate validation results are reported in Section 3.

### 2.6 Linear-quadratic feedback control

For each memory load and routing configuration, the augmented model identified in Eq. (6) is used to synthesize a discrete-time LQR controller. During closed-loop simulation, the full nonlinear spiking network remains the controlled plant. At each 10-ms controller sample, the IPS task state *x*_*k*_ is remeasured from the spiking network, while the two-dimensional actuator state is propagated using the identified actuator dynamics. The feedback state is therefore defined as

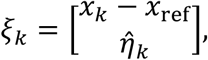

where *ξ*_*k*_ ∈ ℝ^*L+2*^ is the state supplied to the LQR feedback law, *x*_*k*_ − *x*_ref_ is the measured mnemonic task-state error, and 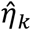 is the estimated actuator-state deviation. The actuator estimate is initialized as 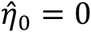 at cue offset and subsequently propagated using Eq. (5b) with the physically applied, non-negative post-cue command.

The feedback gain is computed offline and held fixed for the corresponding load and routing configuration. During the 2500-ms post-cue delay, a new control command is computed every Δ*t*_*c*_ *=* 10 ms and held constant over the following controller interval. The physically applied post-cue command therefore varies with the measured mnemonic state over the delay.

The LQR objective penalizes mnemonic task-state error and control-command magnitude according to the following infinite-horizon quadratic objective

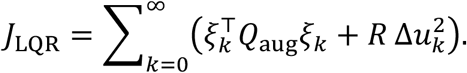

The augmented state-cost matrix is considered as

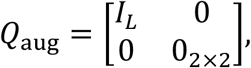

where *I*_*L*_ is the *L × L* identity matrix, that means all *L* task coordinates receive equal state weight, and no direct cost is assigned to the two actuator-state coordinates. The scalar command penalty is fixed at

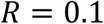

for every memory load and both routing configurations. The feedback gain *K* is obtained from the stabilizing solution *P* of the discrete-time algebraic Riccati equation [50], giving

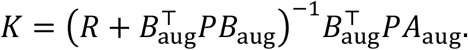

The unconstrained LQR command is

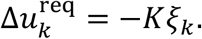

Because the implemented control pathway supplies additional excitatory current to dlPFC, negative commands are not physically applied. The command delivered to the spiking network is therefore

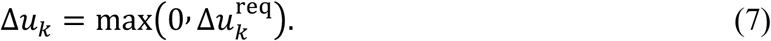

The realized post-cue control effort of each closed-loop trajectory is quantified by the integrated squared physically applied command,

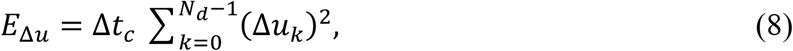

where *N*_*d*_ *=* 250 is the number of controller intervals in the 2500-ms delay and Δ*t*_*c*_ *=* 0.01s. *E*_Δ*u*_ is reported in nA^*2*^ s and serves as the realized model-based post-cue control-effort measure used to examine load-dependent demands on the dlPFC pathway.

To assess the contribution of the temporal structure of state feedback, we also evaluate a mean-matched open-loop condition. For each realization, the mean physically applied LQR command over the delay is first computed as

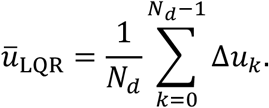

A separate open-loop trajectory then receives 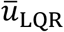 as a constant post-cue current while using the same prefrontal recruitment protocol.

### 2.7 Localized disturbance tests

To assess the contribution of state feedback to robustness, localized suppressive disturbances were applied under cue-aligned selective routing at memory load 5. The disturbance was injected directly into IPS excitatory neurons around the first mnemonic location (36°) using a circular Gaussian spatial profile,

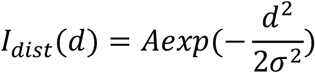

with σ *=* 9.4*°*. Here, *d* denotes the circular angular distance from the disturbance center and *A* denotes the peak suppressive current. Two disturbance conditions were examined: a temporary disturbance with *A =* −0.25 nA applied from 2000 to 2400 ms, and a sustained disturbance with *A =* −0.17 nA applied from 2000 ms to the end of the 3000-ms trial. Closed-loop and mean-matched open-loop trajectories were initialized from the same mnemonic state and received identical stochastic inputs and disturbance currents. In the open-loop condition, the command was fixed at the mean applied command of the corresponding nominal LQR trajectory, whereas the closed-loop command was recomputed every 10 ms from the measured mnemonic state.

### 2.8 Finite-horizon reachability and mnemonic regulation demand

Finite-horizon reachability analysis is used to characterize the local geometry of dlPFC access to the cue-defined mnemonic task space. The analysis is performed on the identified task-actuator model and is therefore interpreted as a model-based description of accessible directions and relative input authority near the identified operating regime, not as a quantitatively accurate 500-ms forecast of the nonlinear spiking network. Although the dlPFC command is scalar, its repeated propagation through the local task-actuator dynamics can produce terminal displacements along more than one task-space direction. We therefore characterize those directions directly from the finite-horizon input-to-task map. For the augmented model in Eq. (6), [51,52]

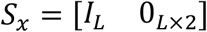

selects the *L* mnemonic task coordinates from the augmented state. Over a horizon of *N* controller intervals, the mapping from a sequence of scalar dlPFC commands to the terminal task-state displacement is

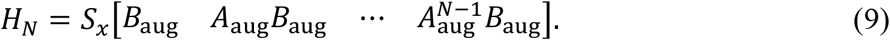

The main analysis uses a 500-ms horizon, corresponding to *N = 50* controller intervals at the 10-ms sampling time. Thus, *H*_*N*_ maps the sequence of 50 scalar dlPFC commands applied over this interval to the resulting terminal displacement in the mnemonic task space. Each column of *H* represents the contribution of a command applied at a particular controller step after propagation through the task–actuator dynamics to the end of the horizon. We characterize the geometry of this finite-horizon input-to-task mapping using the singular-value decomposition

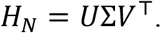

The singular-value decomposition separates this local input-to-task map into orthogonal task-space directions and their associated gains. The left singular vectors, given by the columns of *U*, define orthonormal directions of terminal mnemonic-state displacement, while the corresponding singular values σ_*i*_ quantify finite-horizon input-to-state gain along those directions. For a fixed norm of the command sequence, larger singular values imply a larger attainable terminal displacement along the associated left singular vector. The normalized singular spectrum, σ_*i*_*/*σ_1_, is used as a descriptive measure of anisotropy. Because the squared singular values are the eigenvalues of the corresponding finite-horizon task-space controllability Gramian, we also summarize the concentration of reachability strength by the leading-mode Gramian share and a participation-ratio effective dimension.

To assess the structure of the dominant reachable direction, we compare the leading left singular vector *v* with an equal-item common-strength direction, defined by equal weighting of all mnemonic coordinates, and quantified their alignment using absolute cosine similarity. We additionally report the ratio of the second to the leading Gramian eigenvalue as a measure of separation between the two strongest finite-horizon directions, together with the numerical row rank of the input-to-task map. These quantities were computed separately for each memory load and routing configuration.

To quantify control authority along the most accessible task-space direction, let *v* denote the leading unit-norm left singular vector of *H*_*N*_, with corresponding singular value σ . The minimum squared-input cost associated with a unit terminal displacement along this direction is

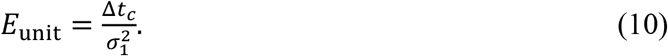

Smaller values of *E*_*unit*_ indicate greater finite-horizon control authority along *v*. This quantity is derived from the local linear reachability model and is used to compare control authority across memory loads and routing configurations. The reachability analysis is based on the unconstrained linear input-to-task map and therefore describes local finite-horizon control authority under signed inputs. By contrast, the command applied in the spiking-network simulations is constrained to be non-negative according to Eq. (7).

Equation (10) provides a normalized measure of directional control authority along the dominant singular mode. For a fixed horizon and unit terminal displacement, it quantifies the minimum unconstrained signed-input energy required by the local linear model to reach that direction. This quantity is conceptually distinct from the realized control effort in Eq. (8), which is computed from the non-negative command applied to the nonlinear spiking network. It is therefore used to characterize local reachability rather than to predict the magnitude of realized LQR effort.

For each singular direction *v*_*i*_, the unit-displacement cost is normalized by that of the dominant mode, *E*_unit,i_*/E*_unit,1_, providing a dimensionless measure of relative directional control authority. Larger ratios indicate weaker finite-horizon access relative to *v*_1_.

We next quantify the mnemonic regulation demand present before additional post-cue control is applied. For the recruitment-only trajectory, the task-space correction required to return the current mnemonic state to its cue-offset reference is defined as

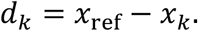

The vector *d*_*k*_ is then decomposed relative to the leading reachable direction *v*:

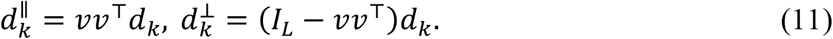

Here, 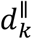 is the component of the mnemonic regulation demand aligned with the leading reachable direction, 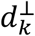 is the component orthogonal to that direction, *v* ∈ ℝ^*L*^ is the leading unit-norm left singular vector of *H*_*N*_, and *I*_*L*_ is the *L × L* identity matrix. The matrix *vv*^*T*^projects *d*_*k*_ onto the one-dimensional subspace spanned by *v*, while *I*_*L*_ − *vv*^*T*^projects it onto the complementary orthogonal subspace. The magnitudes of the two components are summarized over the delay for each independent test realization and compared across memory loads and routing configurations.

The parallel and orthogonal correction magnitudes were normalized by the task-space dimension to support comparison across loads. These quantities describe how mnemonic regulation demand is distributed relative to the dominant reachable direction.

## 3. Results

### 3.1 Local task–actuator model validation

The held-out one-step validation results for all ten load-by-routing models are summarized in Table 1. Across conditions, the identified models reproduced the direction of the measured state changes with consistently high cosine similarity (0.953–0.997), while the corresponding NRMSE values ranged from 19.4% to 35.1%, indicating that the local models captured the direction of state evolution more consistently than its precise magnitude.

**Table 1.** Held-out one-step validation of the local task–actuator model across memory loads and routing configurations. NRMSE quantifies prediction-error magnitude and cosine similarity quantifies directional agreement between measured and predicted one-step responses.

| Memory load | Flat routing |  | Cue-aligned selective routing |  |
| --- | --- | --- | --- | --- |
|  | NRMSE (%) | Cosine similarity | NRMSE (%) | Cosine similarity |
| 3 | 29.2 | 0.968 | 30.4 | 0.953 |
| 4 | 33.5 | 0.972 | 26.1 | 0.967 |
| 5 | 28.4 | 0.987 | 27.0 | 0.978 |
| 6 | 28.5 | 0.997 | 35.1 | 0.970 |
| 7 | 19.4 | 0.991 | 26.0 | 0.988 |

Figure 3 illustrates the cue-aligned selective model at load 5 as a representative case. The predicted trajectories closely followed the measured one-step responses of both the five mnemonic coordinates and the two actuator coordinates, with aggregate NRMSE of 27.0% and cosine similarity of 0.978. Taken together, the validation results reflects the broader validation pattern across loads and routing configurations: the identified task–actuator models reproduce the local directional structure of the measured responses sufficiently well for controller synthesis and geometric analysis, without implying quantitative equivalence to the nonlinear spiking network over extended horizons. In Fig. 3, panels x1-x5 each show one mnemonic task coordinate, followed by the actuator p panel and the actuator a panel; every panel compares the measured one-step response with the corresponding model prediction.

**Fig. 3.**
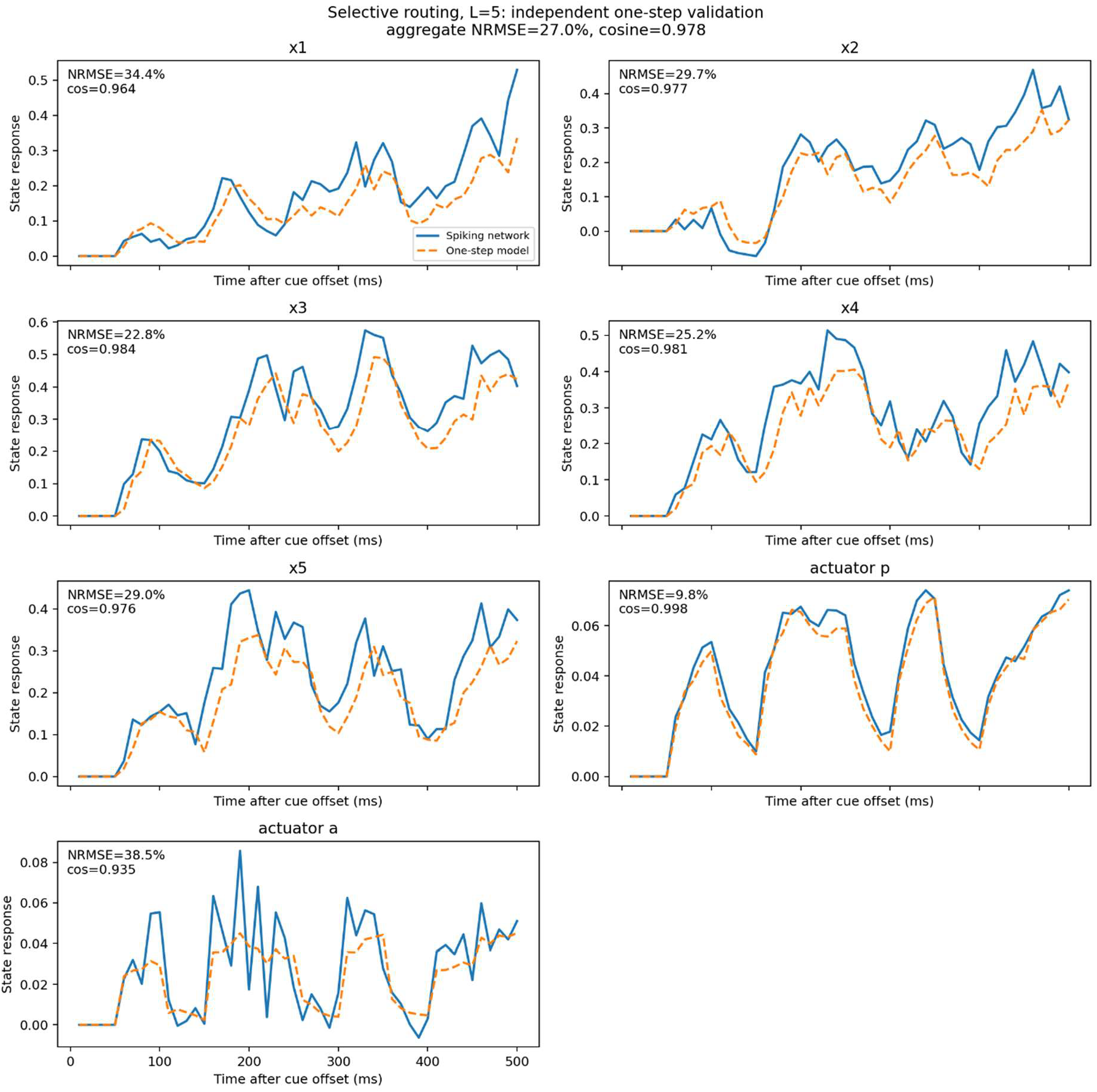
Representative one-step validation of the local task–actuator model at memory load 5 under cue-aligned selective routing. Measured and predicted responses are shown for the five mnemonic and two actuator coordinates; aggregate validation results are reported in Table 1.

### 3.2 Load-dependent working memory retention under flat and cue-aligned selective routing

Before evaluating post-cue control, we first established an Edin-like recruitment-only baseline. A 0.15-nA current pulse was applied to dlPFC excitatory neurons during the first 100 ms of cue presentation to recruit persistent prefrontal activity, after which no additional post-cue control command was applied. Figure 4A compares late-delay retention in the intrinsic IPS network with this recruitment-only flat-routing condition. Prefrontal recruitment increased retention relative to the intrinsic IPS network, reproducing the qualitative capacity-enhancement effect reported by Edin et al. [20] and providing the baseline for the subsequent control comparisons.

**Fig. 4.**
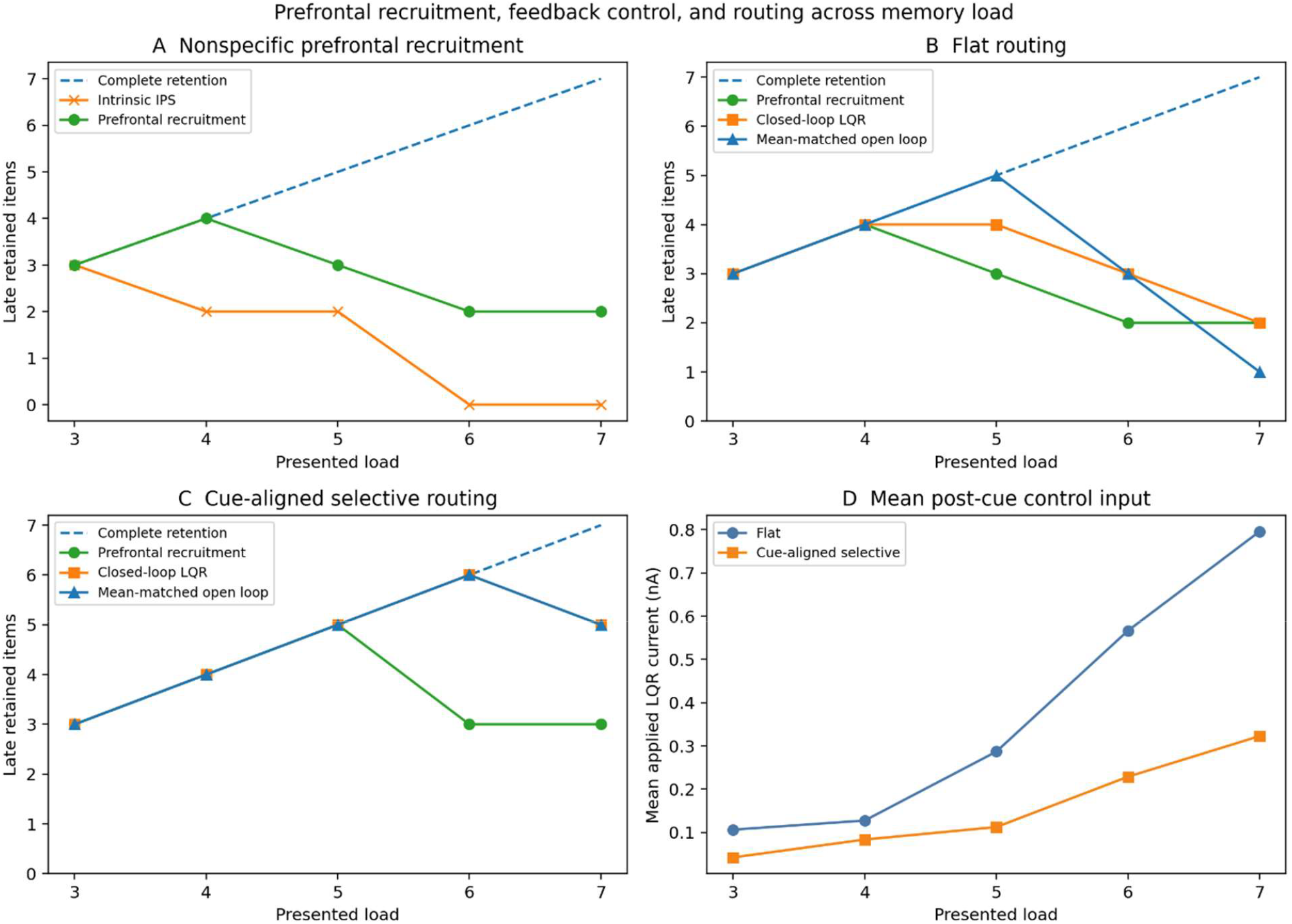
Representative prefrontal recruitment, post-cue control, and routing across memory load. (A) Late retained-item count for the intrinsic IPS network and the Edin-like recruitment-only flat-routing baseline. (B) Late retained-item count under flat routing for recruitment only, closed-loop LQR, and mean-matched open-loop control. (C) Corresponding late retained-item count under cue-aligned selective routing. (D) Mean physically applied post-cue LQR command for flat and selective routing in the same representative realization. The dashed line denotes complete retention; across-realization statistics are reported in the text and Table 2.

Figure 4B and C compare recruitment-only, closed-loop LQR, and mean-matched open-loop control under flat and cue-aligned selective routing, respectively, while Fig. 4D shows the mean physically applied post-cue LQR command for the same representative realization. In this realization, closed-loop LQR retained 3, 4, 4, 3, and 2 items under flat routing and 3, 4, 5, 6, and 5 items under selective routing across loads 3–7, respectively. Across five independent test realizations, complete retention was maintained through load 4 under flat routing and through load 6 under selective routing. At load 5, the mean retained-item count was 3.2 under flat routing and 5.0 under selective routing; at load 6, the corresponding values were 2.6 and 6.0.

**Table 2.** Mean physically applied post-cue LQR command and realized post-cue control effort across memory loads and routing configurations.

| Memory load | Flat mean applied command (nA) | Selective mean applied command (nA) | Flat realized control effort (nA<sup>2</sup> s) | Selective realized control effort (nA<sup>2</sup> s) |
| --- | --- | --- | --- | --- |
| 3 | 0.092 ± 0.016 | 0.034 ± 0.010 | 0.038 ± 0.009 | 0.005 ± 0.002 |
| 4 | 0.141 ± 0.014 | 0.092 ± 0.015 | 0.068 ± 0.012 | 0.031 ± 0.007 |
| 5 | 0.289 ± 0.049 | 0.110 ± 0.012 | 0.267 ± 0.085 | 0.041 ± 0.007 |
| 6 | 0.682 ± 0.135 | 0.216 ± 0.010 | 1.339 ± 0.550 | 0.136 ± 0.012 |
| 7 | 0.911 ± 0.128 | 0.294 ± 0.021 | 2.241 ± 0.596 | 0.260 ± 0.029 |

Neither routing configuration reliably maintained all seven items. Mean-matched open-loop control produced comparable late-delay retention, including complete retention in every selective-routing realization through load 6. Thus, cue-aligned routing shifted the retention boundary and reduced the required post-cue drive, whereas replacing state-dependent feedback with a mean-matched constant command had little effect on nominal retention. The across-realization command and effort trends are quantified in Section 3.4 and Table 2.

Load 6 provides a direct illustration of the routing effect (Fig. 5). Under flat routing, closed-loop IPS activity retained three of six mnemonic representations (Fig. 5A), and the corresponding late-delay firing profile shows the surviving representations under closed-loop LQR and mean-matched open-loop control (Fig. 5B). Figure 5C shows the full-trial dlPFC inputs, including the common recruitment pulse and the post-cue commands: the unconstrained LQR output, the non-negative command actually applied to dlPFC after rectification, and the mean-matched open-loop command, which is held constant at the mean applied LQR level. The unconstrained and physically applied LQR commands overlap whenever the unconstrained output is non-negative; they differ only when negative unconstrained values are clipped to zero. Under cue-aligned selective routing, all six mnemonic representations were maintained (Fig. 5D), as also evident in the late-delay firing profiles (Fig. 5E); the corresponding full-trial dlPFC inputs are shown in Fig. 5F. By comparison, the Edin-like recruitment-only condition retained two of six items (Fig. 5G,H); its input consisted solely of the 0.15-nA, 100-ms recruitment pulse, with no additional post-cue command (Fig. 5I). In this representative realization, selective routing maintained 6/6 items with a mean physically applied post-cue command of 0.229 nA, whereas flat routing retained 3/6 items despite a mean command of 0.566 nA. The mean-matched open-loop trajectories yielded the same retained-item counts. Thus, the representative dynamics mirror the across-realization result: selective routing supported higher retention with substantially less top-down drive. Across-load command and realized-effort statistics are reported in Section 3.4 and Table 2.

**Fig. 5.**
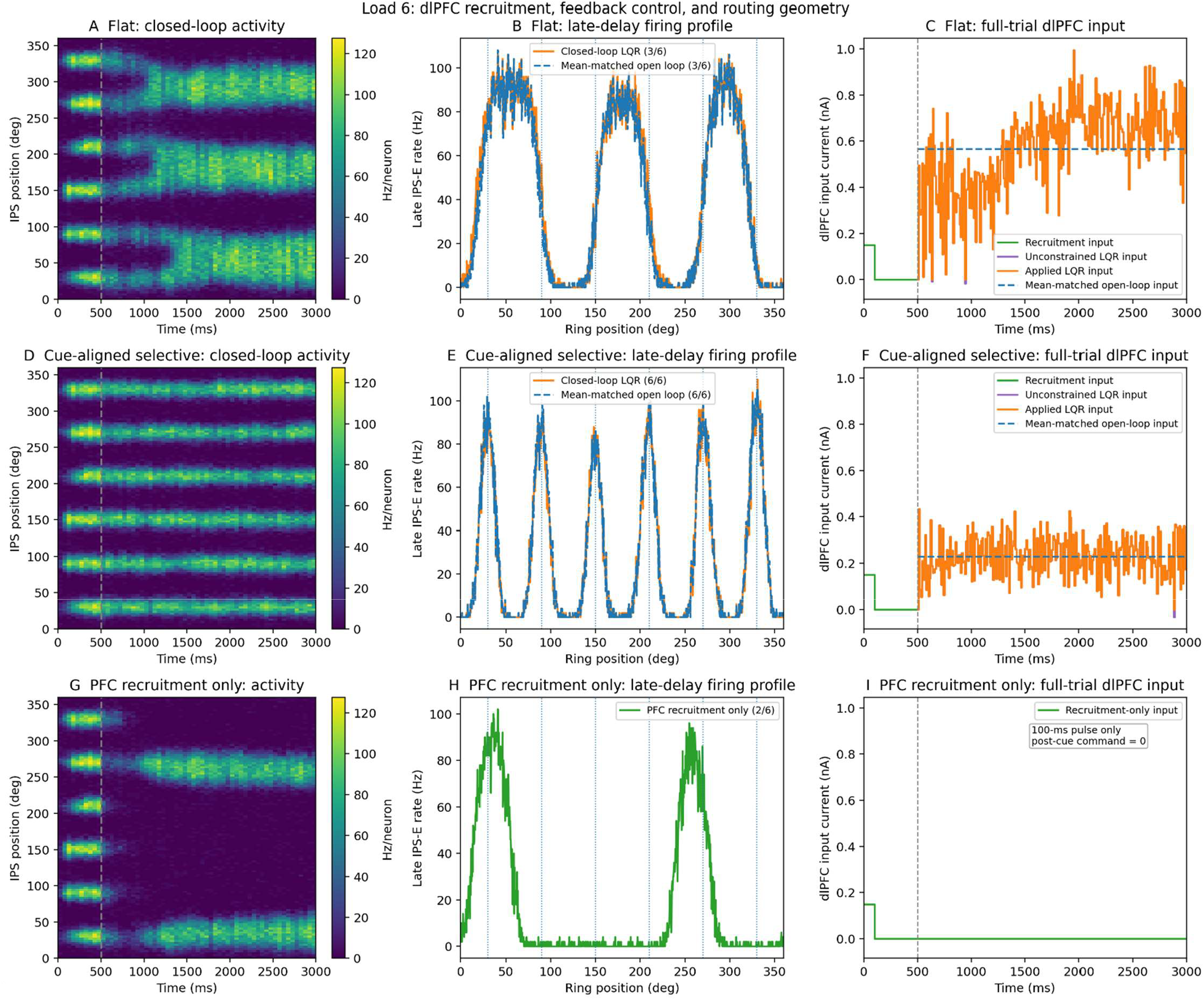
Representative load-6 dynamics under flat, cue-aligned selective, and Edin-like recruitment-only conditions. (A–C) Flat-routing IPS activity, late-delay firing profiles, and dlPFC input. (D–F) Corresponding selective-routing results. (G–I) Recruitment-only activity, late-delay profile, and input. Closed-loop flat and selective routing retained 3/6 and 6/6 items, respectively; recruitment only retained 2/6.

### 3.3 Closed-loop feedback enhances robustness to localized mnemonic disturbance

Because closed-loop LQR and the mean-matched open-loop condition produced similar late-delay retention under nominal conditions, we next tested whether state feedback provided an advantage when the mnemonic state was displaced from its nominal trajectory. At memory load 5 under cue-aligned selective routing, a localized suppressive disturbance was applied to one maintained IPS representation (Fig. 6, Fig. 8). The disturbance waveform was identical across control conditions. In the mean-matched open-loop condition, the post-cue command remained fixed at the mean physically applied LQR command from the corresponding nominal trajectory. In the closed-loop condition, by contrast, the command was recomputed every 10 ms from the measured mnemonic-state deviation relative to the fixed cue-offset reference *x*_*ref*_. Thus, the comparison isolates the benefit of state-dependent adaptation relative to a constant command of comparable nominal magnitude.

**Fig. 6.**
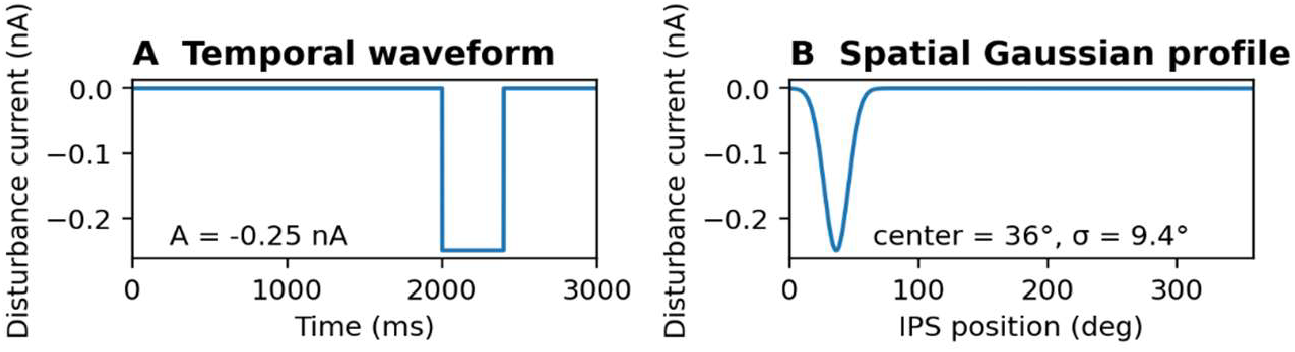
Temporary localized disturbance at memory load 5. A Gaussian suppressive input (−0.25 nA peak) was applied to the first mnemonic location from 2000 to 2400 ms, independently of dlPFC control.

Under the temporary disturbance (Fig. 6), the retained-item count remained at 5/5 under closed-loop LQR (Fig. 7A), whereas the mean-matched open-loop condition transiently dropped to 4/5 before recovering after disturbance offset. The closed-loop command increased during the disturbance (Fig. 7B), consistent with compensatory state-dependent recruitment. The corresponding IPS activity shows that the targeted representation remained preserved under closed-loop control (Fig. 7C), while it was strongly suppressed under open-loop control before re-emerging after the disturbance ended (Fig. 7D).

**Fig. 7.**
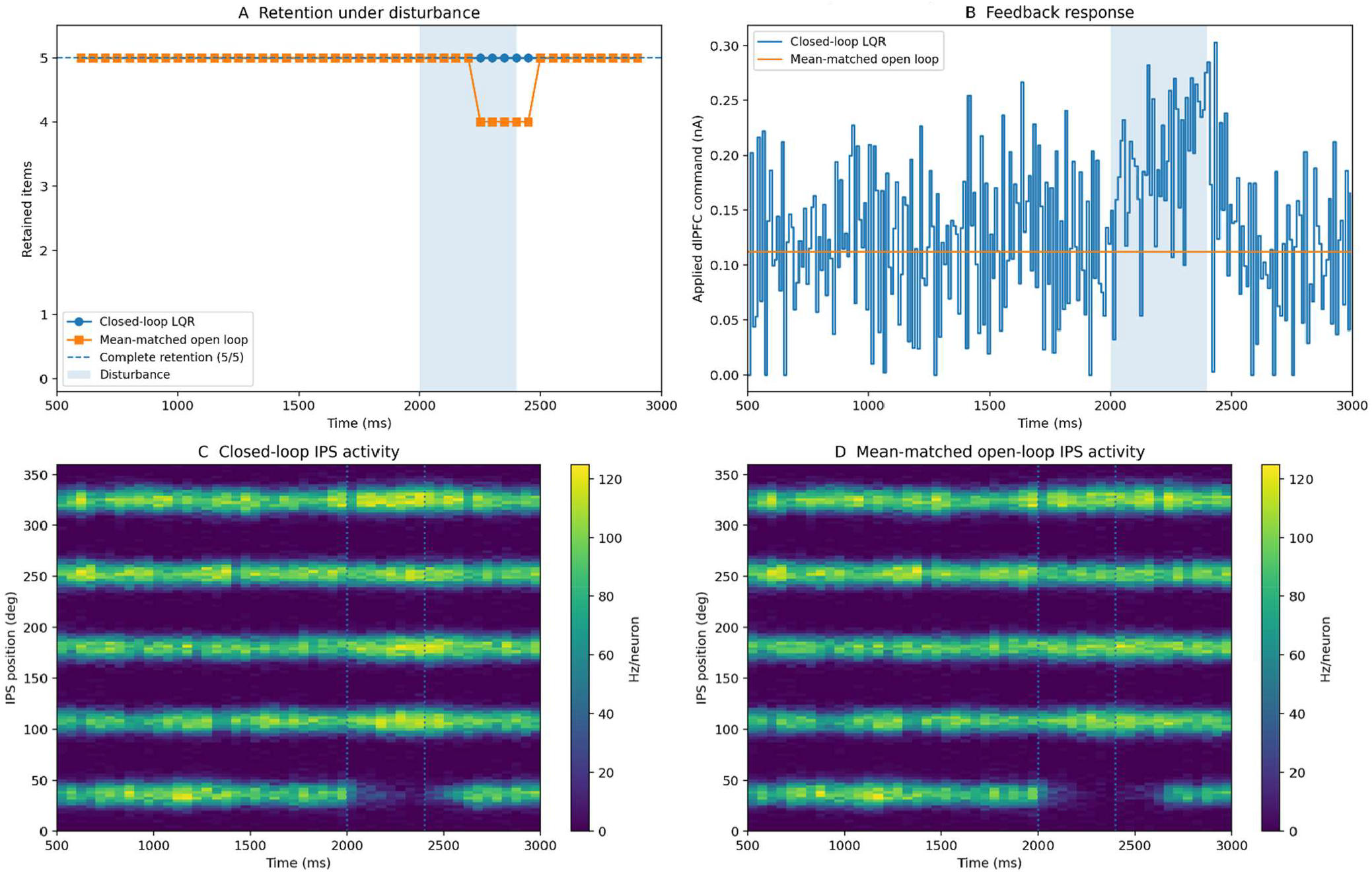
Temporary disturbance rejection at memory load 5. (A) Retained-item count under closed-loop LQR and mean-matched open-loop control. (B) Applied dlPFC command during the disturbance. (C,D) IPS activity under closed-loop and open-loop control. Closed-loop feedback preserves all five representations, whereas open-loop control transiently loses the targeted item.

Importantly, recovery in the open-loop condition does not depend on *x*_*ref*_, because the reference is not used by the open-loop policy. The temporary disturbance strongly suppresses the targeted representation but does not completely extinguish its underlying recurrent activity. Once the suppressive input is removed, this residual mnemonic activity, together with the continuing constant dlPFC drive and recurrent IPS dynamics, allows the weakened bump to recover toward the attractor. In the closed-loop formulation, by contrast, the cue-offset mnemonic state defines the fixed reference *x*_ref_, which is used only to compute the state deviation driving the scalar dlPFC command; the reference itself is never directly imposed on the IPS state.

The sustained disturbance (Fig. 8) produced a stronger separation between the two control strategies. Closed-loop LQR maintained 5/5 retention throughout the disturbance (Fig. 9A), accompanied by an adaptive increase in the applied dlPFC command (Fig. 9B), and preserved the targeted IPS representation (Fig. 9C). In contrast, the mean-matched open-loop condition lacked state-dependent compensation and the targeted representation remained suppressed, yielding 4/5 retained items at the end of the trial (Fig. 9D). Together, these results show that mean-matched tonic prefrontal drive is sufficient for nominal maintenance, whereas state-dependent feedback becomes advantageous when mnemonic activity is displaced from its nominal trajectory.

**Fig. 8.**
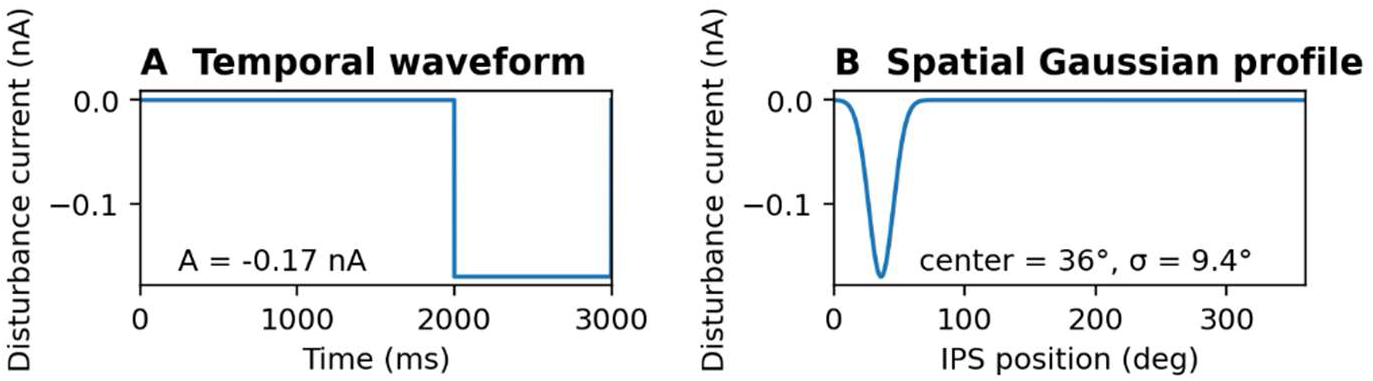
Sustained localized disturbance at memory load 5. A Gaussian suppressive input (−0.17 nA peak) was applied to the first mnemonic location from 2000 ms to trial end, independently of dlPFC control.

**Fig. 9.**
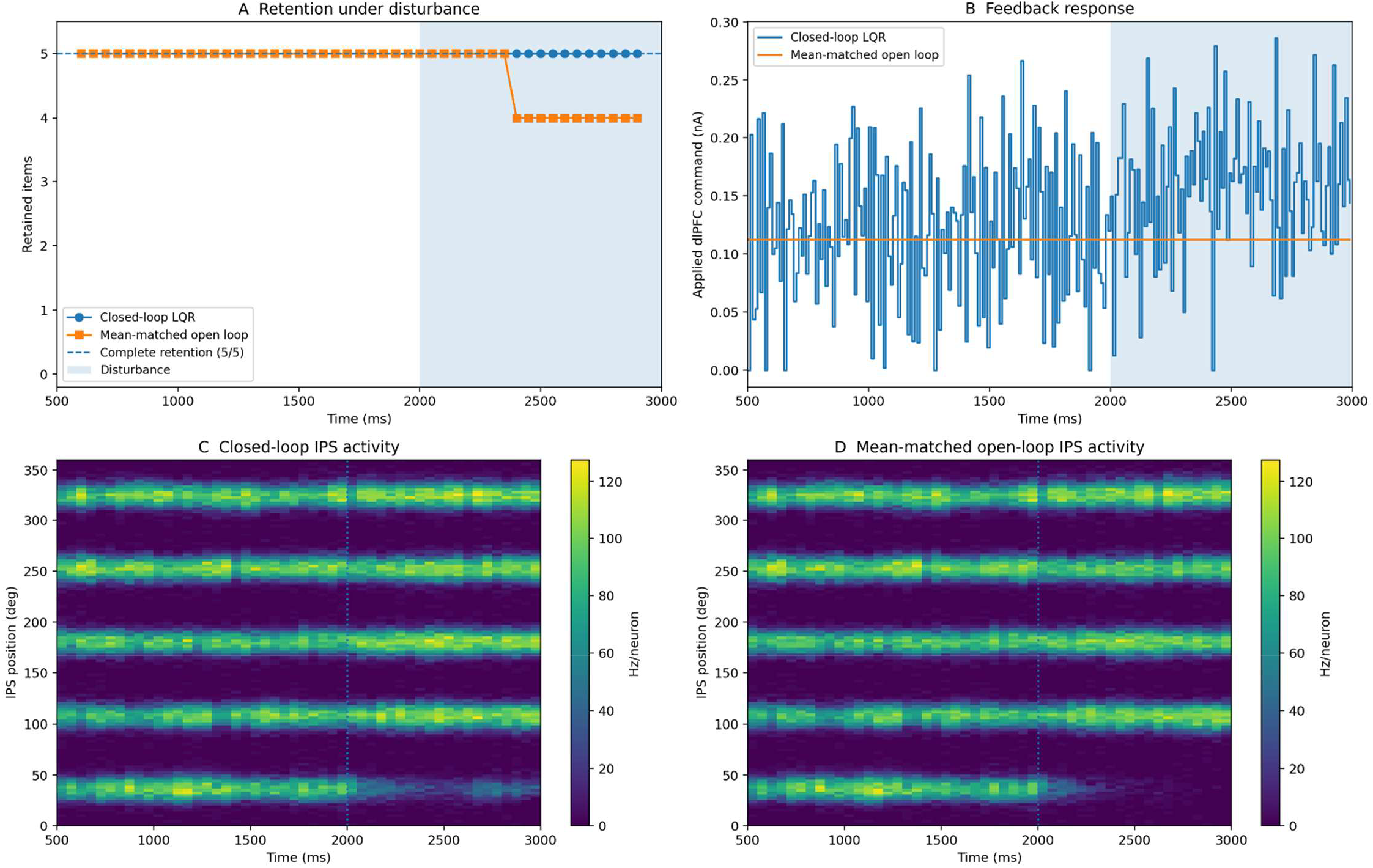
Sustained disturbance rejection at memory load 5. (A) Retained-item count. (B) Physically applied dlPFC command; shading indicates the disturbance interval. (C,D) IPS excitatory activity under closed-loop and mean-atched open-loop control, respectively. Closed-loop LQR maintains 5/5 retention, whereas open-loop control loses the targeted representation and ends with 4/5 retained items.

### 3.4 Realized post-cue control effort increases with memory load

The physically applied post-cue LQR command rose systematically with memory load and was consistently lower under cue-aligned selective routing (Table 2). Values are mean ± SD across five independent test realizations. Realized post-cue control effort is the integrated squared physically applied LQR command defined in Eq. (8); the common prefrontal recruitment pulse is excluded. Table 2 reports the across-realization numerical summary. In comparison, Fig. 4D shows the mean applied command in one representative realization, while Fig. 10A visualizes the across-realization trend in realized control effort.

**Fig. 10.**
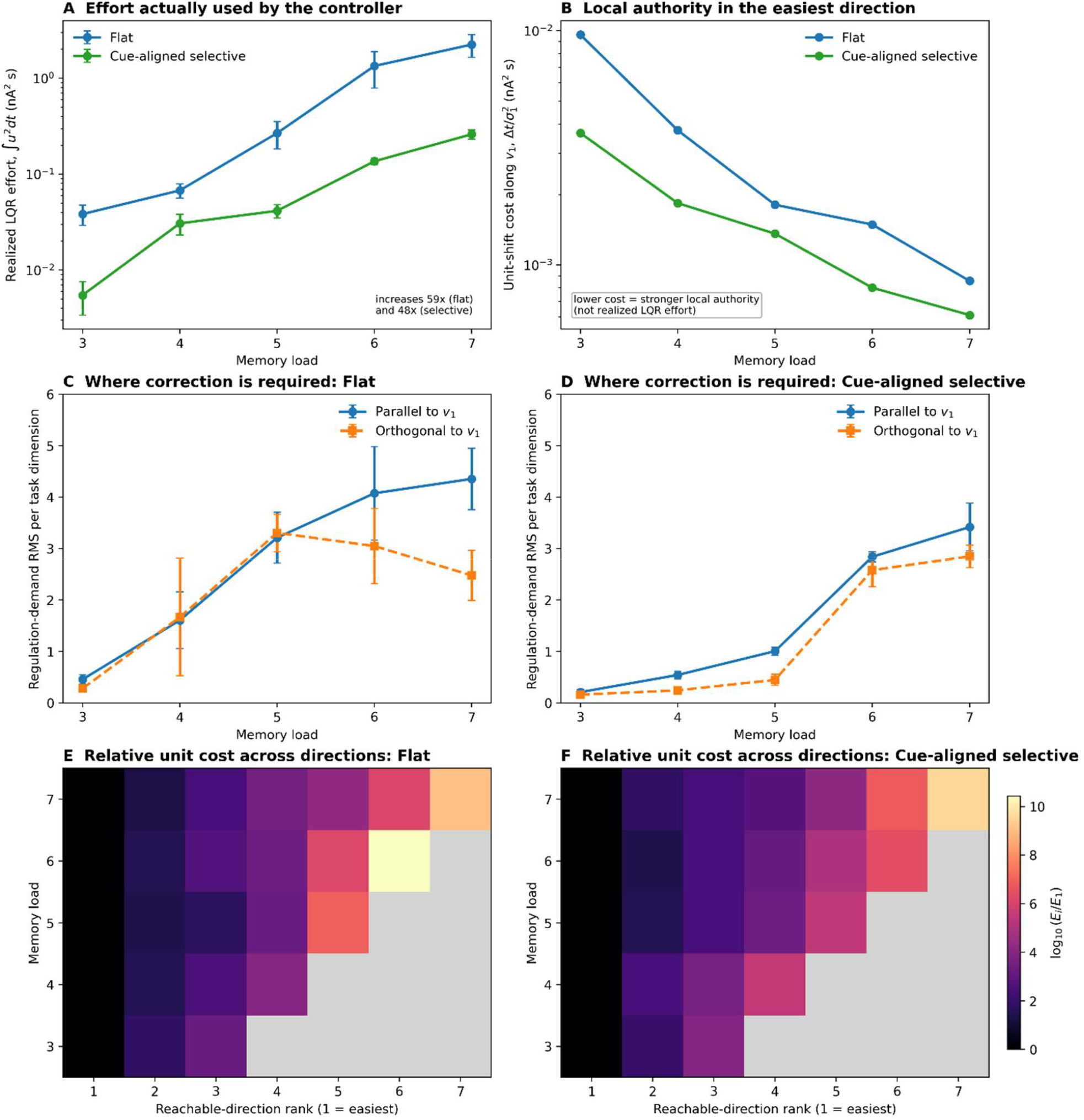
Realized control effort, local reachability, and regulation demand. (A) Integrated squared physically applied LQR command (mean ± SD across five realizations). (B) Unit-displacement cost along the leading direction *v*_1_; lower alues indicate greater local control authority. (C,D) Recruitment-only regulation demand parallel and orthogonal to *v*_1_ under flat and cue-aligned selective routing, respectively. (E,F) Relative unit-displacement cost across singular directions under flat and selective routing, normalized to the leading direction and shown on a log scale.

The corresponding realized control-effort measure showed a stronger load dependence. Integrated squared command increased from 0.038 ± 0.009 to 2.241 ± 0.596 nA^2^ s under flat routing and from 0.005 ± 0.002 to 0.260 ± 0.029 nA^2^ s under selective routing, equivalent to approximately 58-fold and 48-fold increases, respectively. This increase was monotonic across loads in every independent realization for both routing configurations. Selective routing reduced realized effort at every tested load, with the largest absolute separation at high load.

To determine whether the load-dependent increase in realized control effort reflected stronger sustained drive or greater temporal modulation, we decomposed each closed-loop LQR command trajectory into its temporal-mean and zero-mean time-varying components. Because

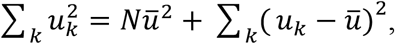

where *u*_*k*_ is the physically applied post-cue command at controller interval *k, N* is the number of controller intervals over the delay period, and 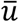 is the temporal mean of the applied command, the two contributions can be quantified separately. The term 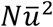 represents the sustained mean-command component, whereas 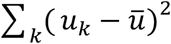 represents the zero-mean time-varying component. Multiplying each term by the controller interval Δ*t* gives its corresponding contribution to the realized control effort defined in Eq. (8). The sustained mean-command component accounted for 55.9%, 73.5%, 79.6%, 90.0%, and 94.0% of total effort under flat routing across loads 3–7, and 51.9%, 69.4%, 73.2%, 85.6%, and 83.3% under selective routing. Thus, the load-dependent increase in realized effort was driven primarily by stronger sustained post-cue dlPFC recruitment, particularly at higher loads, while time-varying modulation generally accounted for a smaller fraction of the total at higher loads.

### 3.5 Finite-horizon reachability geometry and mnemonic regulation demand

Finite-horizon analysis revealed a strongly anisotropic reachability geometry for dlPFC access to the mnemonic task space. Although the 500-ms input-to-task reachability matrices were numerically full row rank, their normalized singular spectra decayed steeply under both routing configurations, leaving only very weak gain along the higher-order directions. Given the scalar untuned actuator, the symmetry of the equally spaced mnemonic locations, and the small gain of these higher-order modes, numerical full rank was not interpreted as evidence for multiple physiologically meaningful independent control channels. Instead, the dominant feature of the local reachability geometry was the concentration of finite-horizon input-to-state gain along a single leading task-space direction.

This concentration was quantified using the finite-horizon task-space controllability Gramian. Across loads 3–7, the ratio of the second to the leading Gramian eigenvalue ranged from 1.47% to 4.87% under flat routing and from 0.33% to 3.51% under cue-aligned selective routing. The leading mode accounted for 94.4–98.5% of the total Gramian trace under flat routing and 96.2– 99.6% under selective routing. Consistent with this strong concentration, the participation-ratio effective dimension remained close to one, ranging from 1.03 to 1.12 for flat routing and from 1.01 to 1.08 for selective routing (Table 3). Thus, finite-horizon dlPFC access in the identified local model was effectively one-dimensional, with substantially weaker access to higher-order task-space directions.

**Table 3.** Finite-horizon task-space controllability metrics across memory loads and routing configurations. The second-to-leading Gramian eigenvalue ratio quantifies separation between the first two controllability modes; the leading-mode share is the fraction of total Gramian trace carried by the first mode; and the effective controllability dimension is the participation-ratio dimension of the normalized Gramian spectrum.

| Memory load | Second/leading eigenvalue ratio (%) |  | Leading-mode share of Gramian trace (%) |  | Effective dimension |  |
| --- | --- | --- | --- | --- | --- | --- |
|  | Flat | Selective | Flat | Selective | Flat | Selective |
| 3 | 1.47 | 1.06 | 98.50 | 98.94 | 1.030 | 1.021 |
| 4 | 3.04 | 0.328 | 96.77 | 99.65 | 1.067 | 1.007 |
| 5 | 4.09 | 2.93 | 94.39 | 96.77 | 1.120 | 1.067 |
| 6 | 3.05 | 3.51 | 96.82 | 96.20 | 1.066 | 1.079 |
| 7 | 4.87 | 1.49 | 94.96 | 97.95 | 1.106 | 1.042 |

Figure 10A shows the strong load-dependent increase in realized LQR effort in the nonlinear closed-loop simulations. Figure 10B provides a complementary model-based measure of local control authority along the dominant reachable direction (*v* ). The unit-displacement cost along this direction decreased with increasing load under both routing configurations. Thus, the rise in realized control effort does not reflect a progressive loss of control authority along the easiest reachable direction.

One possible interpretation is that these two effects reflect different aspects of the mnemonic dynamics. As memory load increases, stronger competition and lateral inhibition make it progressively more difficult to maintain multiple distinct activity bumps. Because the dominant reachable direction is closely aligned with an equal-item common-strength mode, the scalar dlPFC command acts coherently across the active mnemonic coordinates. As load increases, this parallel action can produce a larger aggregate displacement along the normalized common mode, increasing its finite-horizon gain and reducing the corresponding unit-displacement cost. This does not imply that overall mnemonic regulation becomes easier: the correction required relative to the reference grows with load and includes components along much more weakly accessible directions.

Figure 10C shows the mnemonic regulation demand under flat routing. As load increased, the required correction became increasingly concentrated along the dominant reachable direction. At loads 6–7, the parallel component rose sharply while the orthogonal component declined, indicating that high-load regulation was dominated by the common-strength mode rather than by item-specific directions. Thus, under flat routing, increasing load primarily enlarged the mismatch along the direction to which dlPFC has the strongest control authority. Figure 10D shows the corresponding demand under selective routing, where the parallel component increased from 0.21 to 3.42 and the orthogonal component from 0.16 to 2.85. These quantities were computed from recruitment-only trajectories before additional post-cue control and therefore characterize the geometry of the correction required relative to the mnemonic reference, not the allocation of LQR control effort. Across conditions, *v* was closely aligned with the equal-item common-strength direction, with absolute cosine similarity ranging from 0.967 to 0.999.

Figure 10E shows the relative unit-displacement costs of the singular directions under flat routing, normalized to the leading direction, and Fig. 10F shows the corresponding costs under selective routing. The second direction alone required 20.5–68.2 times the leading-direction unit cost under flat routing and 28.5–305.3 times that cost under selective routing, with substantially greater costs along the weaker higher-order directions. Taken together, Fig. 10A–F shows that realized control effort increases strongly with memory load despite preserved local authority along the dominant reachable mode. At the same time, the mnemonic correction required relative to the reference becomes markedly larger and extends into directions that are much more weakly accessible. This pattern is consistent with increasing regulation demand, rather than a progressive loss of authority along the dominant mode, as the more relevant correlate of the rising control requirement. More generally, both routing configurations provide strong access to a common mnemonic mode and much weaker access to independent item-differentiating directions.

## 4. Discussion

The present study used linear control theory to operationalize cognitive effort as formal control demand in a biophysically grounded model of visual working memory, while retaining the nonlinear spiking circuit as the controlled plant. Three main findings emerged. First, realized post-cue control effort increased strongly and monotonically with memory load, with the increase dominated at higher loads by sustained prefrontal drive. Second, the spatial organization of dlPFC→IPS input substantially altered both mnemonic capacity and the control effort required to support it: cue-aligned selective routing supported higher loads with less post-cue drive than spatially nonspecific routing. Third, sustained prefrontal recruitment accounted for much of the nominal retention benefit, whereas state-dependent feedback became particularly valuable when mnemonic activity was displaced from its nominal trajectory. Together, these findings separate the magnitude, spatial organization, and adaptive regulation of prefrontal support and provide a control-theoretic account of why increasing working-memory demand can require progressively greater top-down control.

### 4.1 Control effort and memory load

The central result for the cognitive-effort question was the pronounced increase in realized post-cue control effort with memory load (Section 3.4; Table 2; Fig. 10A). Integrated squared physically applied command increased monotonically across loads in every independent test realization, by approximately 58-fold under flat routing and 48-fold under cue-aligned selective routing. Decomposition of this effort showed that the sustained mean-command component accounted for approximately 52–94% of the total across conditions and became dominant at higher loads. Increasing mnemonic demand was therefore expressed primarily as a requirement for stronger sustained dlPFC support, with comparatively less contribution from moment-to-moment modulation at higher loads. Within the controllosphere framework [1,7], this provides a concrete computational link between increasing working-memory demand and increasing formal control demand. Although the present simulations do not directly measure subjective effort or metabolic expenditure, the pattern provides model-based support for accounts linking sustained cognitive demand to increasing control-related costs [1,5,7]. At a different spatial scale, human network-control studies likewise show that working-memory demand is associated with systematic changes in state stability, control energy, and functional controllability [35,36].

The finite-horizon analysis provides a complementary geometric perspective on these control demands. Across loads and routing configurations, most model-based reachability strength was concentrated along a single direction closely aligned with an equal-item common-strength mode, whereas higher-order item-differentiating directions were substantially more weakly accessible. Importantly, the increase in realized effort with memory load was not accompanied by a progressive loss of local authority along this dominant mode (Fig. 10B). The rising effort therefore cannot be explained simply by deterioration of control authority in the easiest reachable direction. The lower dominant-mode cost at higher load is therefore best interpreted as greater aggregate authority over a collective common mode, not as reduced overall control demand. Instead, the mnemonic correction required relative to the reference increased substantially across the load range, both along the dominant mode and in more weakly accessible directions (Fig. 10C–F). These observations do not establish a causal decomposition of realized LQR effort, because the reachability measures are derived from the local unconstrained model whereas realized effort is measured from the non-negative command applied to the nonlinear spiking network. They are nevertheless consistent with increasing regulation demand within a strongly anisotropic control geometry being more relevant to the load-dependent rise in control requirement than a loss of authority along the dominant mode.

A qualitatively similar concentration of controllability strength in one dominant Gramian mode was reported for the Stroop control model [7]. Although the two models implement very different cognitive operations, this convergence raises the broader hypothesis that prefrontal control may often act through a comparatively low-dimensional set of effective control directions. If so, task demands that extend into weakly accessible dimensions could require disproportionately greater control input. Testing this possibility will require systematic comparisons across tasks, circuit models, and control architectures; the present results establish the relevant geometry in the working-memory model but do not by themselves demonstrate a general property of biological prefrontal control.

### 4.2 Spatial routing

The routing manipulation further shows that control demand depends on how prefrontal influence is delivered to the mnemonic circuit, not only on its overall magnitude. With matched mean dlPFC→IPS projection strength, spatially nonspecific routing reliably supported complete retention through four items, whereas cue-aligned selective routing supported complete retention through six and required substantially less post-cue command and realized effort. Thus, identical population-average projection strength can produce markedly different functional outcomes depending on its spatial organization. This result extends the original Edin et al. [20] account by showing that the geometry of top-down access can shape both effective working-memory capacity and the control cost required to sustain it. More broadly, it suggests that the efficiency of frontoparietal regulation may depend critically on how well the available control pathway is aligned with the task-relevant neural representation. Furthermore, the six-item capacity obtained with cue-aligned selective routing should be interpreted as a property of the particular IPS operating regime and control architecture examined here, not as a proposed universal capacity limit. The selective profile was prescribed from the cued locations in order to isolate the computational consequences of spatially aligned frontoparietal access while holding mean projection strength constant. How comparable routing is established, maintained, or flexibly reconfigured in biological frontoparietal circuits remains an open question. Given well-established individual variability in visual working-memory capacity and its association with attentional control [14,16], variation in effective frontoparietal routing provides one possible mechanism worth testing, but the present model does not establish such a relationship.

### 4.3 Open-versus closed-loop control

The comparison with mean-matched open-loop stimulation distinguishes sustained support from adaptive feedback. Under nominal conditions, mean-matched open-loop drive produced retention outcomes close to those obtained with closed-loop LQR, indicating that the sustained level of prefrontal recruitment and the routing geometry account for much of the capacity enhancement observed in the undisturbed network. Feedback became functionally important when the mnemonic state was displaced by an external disturbance. In those conditions, the closed-loop controller adjusted dlPFC drive as the measured mnemonic state departed from its reference, whereas the open-loop command remained fixed at its nominal mean level. This adaptive response preserved the targeted representation more robustly under both temporary and sustained disturbances. The results therefore suggest complementary roles for tonic and feedback-dependent prefrontal support: sustained recruitment can stabilize nominal mnemonic maintenance, whereas state-dependent regulation provides additional robustness when the maintained state is perturbed.

The role of the mnemonic reference also clarifies how the LQR formulation should be interpreted. In the closed-loop implementation, *x*_*ref*_ is explicitly retained as the fixed cue-offset target required for reference-based regulation, and the measured deviation from this target determines the scalar dlPFC command. The reference itself is not imposed on the IPS state and is not used to reconstruct a suppressed mnemonic representation directly. Accordingly, the present use of LQR should be understood as a normative computational device for defining control demand and generating a state-dependent benchmark policy, not as a claim that biological dlPFC implements the same algorithm or represents the reference in the same explicit coordinate format. The recovery of the temporarily suppressed representation under mean-matched open-loop control, which never uses *x*_*ref*_, further shows that mnemonic recovery can arise from the remaining recurrent IPS state together with sustained prefrontal drive rather than from direct reinstatement of the reference pattern.

### 4.4 Methodological issues

Methodologically, the study uses the identified task–actuator model only where a local reduced representation is required. Held-out validation showed consistently high directional agreement with one-step responses of the spiking network, alongside moderate prediction-error magnitude (Section 3.1; Table 1; Fig. 3). The reduced model is therefore used for controller synthesis and local control-geometry analysis, while the nonlinear spiking network remains the controlled plant and the IPS mnemonic state is remeasured directly at every 10-ms controller update. This equation-free strategy was adopted after an initial conventional mean-field reduction did not preserve the cue-specific local dynamics and spatial bump structure with sufficient fidelity for the present controller-identification problem. The result should not be interpreted as a general limitation of mean-field descriptions; instead, direct identification provided a practical way to retain the microscopic circuit dynamics relevant to the local mnemonic operating state while still obtaining a tractable model for control analysis.

Several considerations delimit the scope of the present conclusions. First, the identified task– actuator models are local approximations around the mnemonic operating regime. Their validation supports their use for one-step response characterization, controller synthesis, and local control-geometry analysis, but not as globally accurate long-horizon surrogates of the nonlinear spiking circuit. This limitation is partly mitigated in the closed-loop simulations because the IPS task state is remeasured directly from the spiking network at every controller update rather than propagated through the reduced model. Second, the cue-aligned routing profile is prescribed from the cued locations and is used as a controlled manipulation of frontoparietal access. The present results therefore establish the computational consequences of spatially aligned routing, but not the biological mechanism by which such routing would be generated or adapted. Likewise, the numerical capacity values depend on the operating regime and parameters of the IPS model and should not be interpreted as universal working-memory limits. Third, realized control effort is defined by the integrated squared non-negative command applied within the specified LQR formulation. Its magnitude therefore depends on the regulation objective, command constraint, and fixed cue-offset reference, and should be interpreted as a formal model-based measure of control demand rather than as a direct measure of subjective effort or metabolic expenditure. The finite-horizon reachability measures provide complementary information about local control geometry; quantitative differences among the weakest higher-order directions are interpreted cautiously because they rely on repeated propagation of the identified local model. Finally, the disturbance analysis was designed as a targeted test of feedback-dependent robustness and considered localized perturbations at one load under selective routing. It therefore demonstrates a robustness advantage of adaptive feedback under the tested conditions rather than an exhaustive characterization of disturbance rejection. In addition, closed-loop and open-loop conditions were mean-matched under nominal operation rather than constrained to equal total input after disturbance onset. Extending these analyses across loads, disturbance classes, and alternative control objectives would provide a broader characterization of feedback robustness.

## 5. Conclusion

This study provides a control-theoretic account of cognitive effort in working memory by quantifying the prefrontal control required to regulate a nonlinear spiking mnemonic circuit. Realized post-cue control effort increased strongly with memory load and was dominated at higher loads by sustained prefrontal recruitment. The spatial organization of that input was equally consequential: cue-aligned selective routing supported higher mnemonic loads with substantially less control effort than spatially nonspecific routing. Mean-matched open-loop stimulation reproduced much of the nominal retention benefit, whereas state-dependent feedback provided additional robustness when mnemonic activity was perturbed. Finite-horizon analysis further revealed a strongly anisotropic, effectively low-dimensional control geometry in which regulation demand increased with load despite preserved authority along the dominant reachable mode. Together, these results suggest that increasing working-memory demand is associated with greater sustained prefrontal control requirement, while the geometry of frontoparietal access determines how efficiently that control can support mnemonic representations.

## Notes

### Competing Interest Statement

The authors have declared no competing interest.

